# Ca^2+^-Dependent Regulation by the Cyclic AMP Pathway of Primary Cilium Length in LLC-PK1 Renal Epithelial Cells

**DOI:** 10.1101/2020.05.11.088336

**Authors:** Noelia Scarinci, Paula L. Perez, María del Rocío Cantero, Horacio F. Cantiello

## Abstract

The primary cilium is a sensory organelle projecting from the apical surface of renal epithelial cells. Dysfunctional cilia have been linked to a number of genetic diseases known as ciliopathies, which include autosomal dominant polycystic kidney disease (ADPKD). Previous studies have determined that renal epithelial primary cilia express both the polycystin-2 (PC2, TRPP2) channel and the type-2 vasopressin receptor (V2R), coupled to local cAMP production. However, little is known as to how Ca^2+^ and cAMP signals lead to changes in the length of the primary cilium. Here, we explored how cAMP signals regulate the length of the primary cilium in wild type LLC-PK1 renal epithelial cells. Primary cilia length was determined by immunocytochemical labeling of the ciliary axoneme. Treatment of cells with the cAMP analog 8-Br-cAMP (1 mM) in normal external Ca^2+^ (1.2 mM) produced a 25.3% increase (p < 0.0001) in the length of the primary cilium, a phenomenon also observed in cells exposed to high external Ca^2+^ (6.2 mM). However, exposure of cells to vasopressin (AVP, 10 μM), which also increases cAMP in primary cilia of LLC-PK1 cells, mimicked the effect of 8-Br-cAMP in normal, but not in high Ca^2+^. Further, specific gene silencing of PC2 expression further increased primary cilium length after 8-Br-cAMP treatment, in normal, but not high Ca^2+^. The encompassed data indicate a crosstalk between the cAMP and Ca^2+^ signals to modulate the length of the primary cilium, in a phenomenon that implicates the expression of PC2.

**Significance Statement:** Morphological changes in primary cilia have been linked to genetic disorders, including autosomal dominant polycystic kidney disease (ADPKD), a major cause of kidney disease. Both cAMP and Ca^2+^ are universal second messengers that regulate polycystin-2 (PC2, TRPP2), a Ca^2+^ permeable non-selective cation channel implicated in ADPKD, and expressed in the primary cilium of renal epithelial cells. Despite current interest, little is known as to how second messenger systems and how aberrant regulation of PC2 may link primary cilium structure with cyst formation in ADPKD. Here we determined that both the cAMP analog 8-Br-cAMP and vasopressin increase the length of the primary cilium in renal epithelial cells. However, this phenomenon depends of external Ca^2+^ and *PKD2* gene silencing. Proper cAMP signaling may be essential in the control of the primary cilium of renal epithelial cells, and the onset of cyst formation in ADPKD.

## Introduction

The primary cilium of renal epithelial cells (Wheatley 1995) is a non-motile organelle that has been linked to mechanical responses serving as a flow sensor (Praetorius 2001, 2003, Nauli 2003, Pazour 2003). It is now accepted that a vast majority of mutated proteins linked to renal cystic diseases can either be localized to the primary cilium or its associated structures such as the basal body, centrosomes or the ciliary transition zone (Hildebrandt 2005). There are more than a hundred known ciliopathies (Badano 2006, Fliegauf 2007, Davis 2012), including syndromes such as Bardet-Biedl (BBS), Joubert (JBTS), Meckel-Gruber (MKS), Alström, Senior-Löken, and Oro-facial-digital type 1 (OFD1), and diseases such as autosomal dominant polycystic kidney disease (ADPKD) and nephronophthisis (NPHP) (Wheatley 1995, Hildebrandt 2007, Davis 2012). ADPKD in particular, is largely caused by mutations in either one of two genes, *PKD1* or *PKD2* that cause the development of epithelial-lined cysts in various organs, including the kidney, liver and pancreas (Hopp 2012, Loftus 2013). The protein products of *PKD1* and *PKD2*, the transmembrane proteins polycystin-1 (PC1) and polycystin-2 (PC2), respectively, have been localized to primary cilia, where they form a functional receptor/ion channel complex (Ong 2013, Ma 2017), whose dysfunction has been associated with renal cysts (Yoder 2002). It remains to be determined, however, how the dysfunctional cilia hypothesis can explain the highly variable renal phenotypic spectrum seen in different diseases (Loftus 2013).

Changes in the length of primary cilia are known to occur in tissues and cells grown under various physiological and pathological conditions (Pazour 2000, Smith 2006, Verghese 2008, Ou 2009). The length of primary cilia in renal cells increases in both mouse kidneys with ischemic injury and human kidney transplants that suffer acute tubular necrosis (Verghese 2009, Verghese 2008). Lithium is a potent pharmacological agent, which has a profound effect in the treatment of bipolar affective disorder and mania (Machado-Vieira 2009). Miyoshi et al. (2009) observed that chronic Li^+^ administration to mice changed the length of primary cilia in the dorsal and ventral striatum of the brain. Primary cilium elongation by Li^+^ was also observed in cultured cells, including neurons and fibroblasts (Miyoshi 2009), suggesting a more general mechanism of action. Abnormally long primary cilia have also been associated with juvenile cystic kidney disease (Smith 2006). Conversely, a mouse model with partial loss of the ciliary protein polaris, shows abnormally short renal primary cilia, associated with animal mortality in autosomal recessive polycystic kidney disease (ARPKD) (Pazour 2000). A recent study showed that maneuvers that block PC2, including Amiloride and Li^+^, elicited the elongation of primary cilia in wild type LLC-PK1 renal epithelial cells, a phenomenon that was mimicked by siRNA gene silencing of PKD2 (Perez 2020). Conversely, high external Ca^2+^, which activates the calcium-sensing receptor (CaSR), induced a strong reduction in ciliary length (Perez 2020). Thus, both Ca^2+^ signals and the PC2 channel, also present in primary cilia (Raychowdhury 2005), seemed implicated in the regulation of renal epithelia primary cilia. However, ciliary length appears to be normal in cells and organisms carrying *PKD1* and *PKD2* mutations, implying defect(s) in function rather than structure (Ong 2013).

In the present study, we evaluated the effect(s) of the cAMP pathway, and/or high external Ca^2+^, that modulates both PC2 expression and activity in renal epithelial cells (Dai 2017), on primary cilium length in the LLC-PK1renal epithelial cell model. The results indicated that both the cAMP analog 8-Br-cAMP and vasopressin lengthened primary cilia, with differential effects associated with the external Ca^2+^ concentration. Opposite effects were also observed by inhibition of *PKD2* gene expression. Thus, activation of the cAMP pathway may be linked to structural changes in the primary cilium, which largely depended upon external Ca^2+^ and the presence of PC2, which may be a key element in the control of the primary cilium in renal epithelial cells. The present results may help explain the initial events of cyst formation in the onset of ADPKD.

## Methods

### Cell culture and immunochemistry

Wild-type LLC-PK1 renal cells were cultured as previously described (Raychowdhury 2005) in Dulbecco’s modified Eagle’s medium (DMEM) supplemented with 3% fetal bovine serum (FBS), without antibiotics. Cells were seeded onto glass coverslips, and grown at 37°C, in a humidified atmosphere with 5% CO_2_ to reach full confluence in two-to-three weeks in culture. Confluent monolayers were used for immunocytochemical studies to assess the length of the primary cilium. Briefly, confluent cell monolayers were overnight exposed to the various experimental conditions, then rinsed twice with phosphate-buffered saline (PBS), and fixed for 10 min in a freshly prepared solution containing para-formaldehyde (4%) and sucrose (2%). Following fixation, cells were washed three times with PBS, blocked for 30 min with BSA (1%) in PBS, and incubated for 60 min with anti-acetylated-α-tubulin antibody (Santa Cruz Biotechnology) to identify primary cilia (Perez 2020), and an FITC-goat anti-mouse secondary antibody (#626511, Invitrogen, Thermo Fisher Scientific, CA, USA). Cells were also counter-stained with DAPI to locate cell nuclei and mounted with Vectashield mounting medium (Vector Laboratories, Burlingame, CA). Each sample was then viewed under an Olympus IX71 inverted microscope connected to a digital CCD camera C4742-80-12AG (Hamamatsu Photonics KK, Bridgewater, NJ). Images were collected and analyzed with the IPLab Spectrum acquisition and analysis software (Scanalytics, Viena, VA), running on a Dell-NEC personal computer.

### Reagents

Unless otherwise stated, chemical reagents, including CaCl_2_, 8-Br-cAMP, and arginine-vasopressin (AVP) were obtained from Sigma-Aldrich (St. Louis, MO, USA), and diluted at their final concentrations as indicated.

### PKD2 gene silencing

Silencing of *PKD2* gene expression in cultured LLC-PK1 cells was conducted using the small interfering RNA technique, as recently reported (Dai 2017). Briefly, two 21-nt PKD2-specific synthetic siRNAs, one of which was a fluorescent (fluorescein) probe, were synthesized by Invitrogen (Buenos Aires, Argentina). A 19-nt irrelevant sequence was also synthesized as scrambled control (Ir-siRNA, Irss). All constructs bore 3’-dTdT overhangs, as originally reported (Wang 2011). The siRNAs sense sequences were as follows; (siPKD2, P1ss) GCUCCAGUGUGUACUACUACA, starting at 906 in exon 3 of the porcine PKD2 gene, and (Ir-siRNA) UUCUCCGAACGUGUCACGU, as scrambled control. siRNA transfection was conducted as follows (Dai 2017): previously confluent cell cultures were trypsinized and grown to 70% confluence in 24-well tissue culture plates containing DMEM supplemented with 3% FBS at 37°C in a 5% CO_2_ atmosphere. Transfection was performed with Lipofectamine 2000 (Invitrogen). Tubes were added either scrambled (Irss, 10 µl) or antisense (P1ss, 10 µl) RNA with Optimum medium (100 μl) in the absence or presence of Lipofectamine (2 μl). Briefly, the tubes were incubated for 5 min at room temperature and then mixed with either Irss or P1ss for another 20 min (200 μl total volume). Cell incubation was conducted by medium change with a mixture of fresh medium (800 μl, DMEM, and 200 μl of the transfection mixture). The total transfection time was 48 h. Silencing efficiency was confirmed by Western blot technique, as previously reported (Dai 2017).

### Measurement of the length of primary cilia in LLC-PK1 cells

The length of extended primary cilia on the fixed confluent monolayer was manually measured by image analysis with the program ImageJ (NIH, Bethesda, MD, USA), as recently described (Perez 2020), and validated by previous studies (Besschetnova 2010, Miyoshi 2009, Ou 2009, Sipos 2018). Briefly, FITC-labeled fluorescent cilia were traced with the “Freehand Line” tool from ImageJ, and measured to obtain its length in pixels (for details, see Perez 2020). The results were then converted into μm with a Neubauer chamber (Hausser Scientific, Horsham, PA, USA). Because the numerical length parameter of primary cilia data followed non-normal distributions under all experimental conditions tested, values were pre-processed by Box-Cox transformation (Box 1964), to aid in a later analysis by parametric statistical tests that require Normal distributions to compare means and SEs (Perez 2020). The potential transformation was used, which is defined as a continuous function that varies with respect to the lambda (λ) power (Kutner 2004). The STATISTICA Version 8.0 program was used to implement these transformations and obtain a “Transformed Length”, where the maximum likelihood approach was used to search for an appropriate λ to minimize the error function. The Box-Cox transformation (Box 1964) was used in the present study as recently reported (Perez 2020), following:

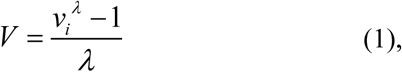

where *V* is the result variable of the transformation, *v*_*i*_ represents the variable to be transformed, and *λ* is the transformation parameter. The transformed distribution histograms showed a remarkable change in their symmetry, approaching Gaussian distributions in all cases (Figs. 1A - 4A). Following data transformation, corrected mean ± standard error values were obtained for each experimental condition and comparisons between groups were conducted by *“t”* Test analysis or one-way ANOVA as appropriate, accepting statistical significance at p < 0.05 (Snedecor 1973).

**Fig. 1:**
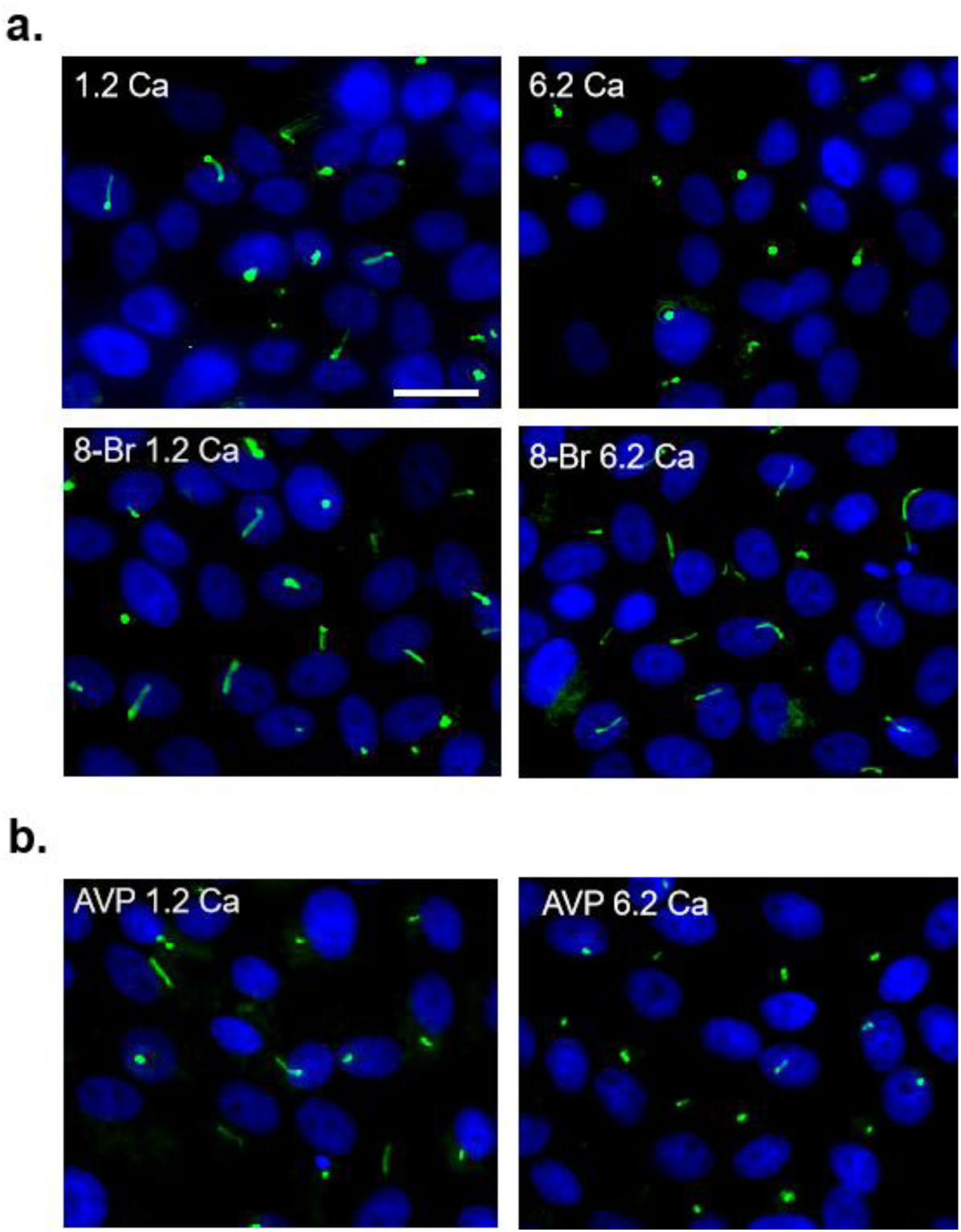
Effect of 8-Br-cAMP, AVP and high Ca^2+^ on the length of primary cilia from LLC-PK1 cells. Confluent monolayers of wild-type LLC-PK1 cells were labeled with an anti-acetylated α-tubulin antibody (Green), and DAPI (Blue). The merged images are shown. Primary cilia were identified (x40) and measured with the IPLab software. **a**. Cells were incubated either in normal (1.2 mM) or high (6.2 mM) external Ca^2+^. Images show a decrease on primary cilium length in high Ca^2+^ with respect to normal Ca^2+^. In the presence 8-Br-cAMP primary cilia were longer in both conditions. **b**. Cells were also exposed to AVP (10 µM) in the presence or absence of high external Ca^2+^. In high external Ca^2+^ (6.2 mM) AVP did not change primary cilium length.

## Results

### Effect of 8-Br-cAMP and high Ca^2+^ on the length of primary cilia from LLC-PK1 cells

To evaluate the effect of the cAMP signaling pathway on the length of the primary cilium of wild type LLC-PK1 cells, confluent monolayers were incubated for 18 h in DMEM medium supplemented with the cAMP analog 8-Br-cAMP (1 mM). Cells were then fixed, and stained with anti-acetylated α-tubulin antibody to measure the length of the primary cilium, as indicated in the Methods section. In the presence of normal external Ca^2+^ (1.2 mM), 8-Br-cAMP-treated cells showed a statistically significant 25.3% increase in primary cilium length as compared to their controls (4.38 ± 0.08 µm, n = 314 vs. 5.49 ± 0.08 µm, n = 235, respectively, p < 0.0001, Fig. 1a, Table 1). Exposure to high Ca^2+^ (6.2 mM) in the absence of 8-Br-cAMP produced a 10.5% decrease in primary cilium length with respect to normal Ca^2+^ (3.92 ± 0.1 μm, n = 242 vs. 4.38 ± 0.08 μm, n = 314, respectively, p < 0.05, Fig. 1a), as recently reported (Perez 2020). The effect of 8-Br-cAMP was then evaluated in the presence of high external Ca^2+^ (6.2 mM), which rendered a cilium length of 5.37 ± 0.12 μm, n = 136. Therefore, under these conditions, 8-Br-cAMP lengthened primary cilium by 1.45 μm, which represented a 37% increase respect to the control condition in high Ca^2+^ (p < 0.0001, Fig. 1a).

**Table 1.** Effect of Ca^2+^ and 8-Br-cAMP on the length of primary cilia

| | 1.2 $\text{Ca}^{2+}$, $\mu\text{m}$ | 6.2 $\text{Ca}^{2+}$, $\mu\text{m}$ | $\Delta$, $\mu\text{m}$ | % |
| --- | --- | --- | --- | --- |
| **Ctrl, $\mu\text{m}$** | $4.38 \pm 0.08$<br>n = 314 | $3.92 \pm 0.11$<br>n = 242 | **-0.46**<br>($p = 0.0006$) | **-10.5** |
| **+ 8-Br-cAMP, $\mu\text{m}$** | $5.49 \pm 0.08$<br>n = 235 | $5.37 \pm 0.12$<br>n = 136 | **-0.12**<br>($p = 0.3894$) | **-2.2** |
| **$\Delta$, $\mu\text{m}$** | **1.11**<br>($p < 0.0001$) | **1.45**<br>($p < 0.0001$) | **0.34** | |
| **%** | **+25.3** | **+37.0** |  |  |

Summarized data shown in Table 1 indicate that 8-Br-cAMP treatment had a stimulatory effect on ciliary length, which was comparatively larger in high external Ca^2+^ (37% vs. 25.3%, Fig. 2a) because of the shortening effect of the ion. The 8-Br-cAMP and external Ca^2+^ effects were independent of each other, and strictly additive, such that a stimulatory effect in length was observed by the cAMP maneuver, and an inhibitory effect by Ca^2+^, which rendered a combined 22.6% (∼1 μm) increase in length by both maneuvers (Table 1).

**Fig. 2:**
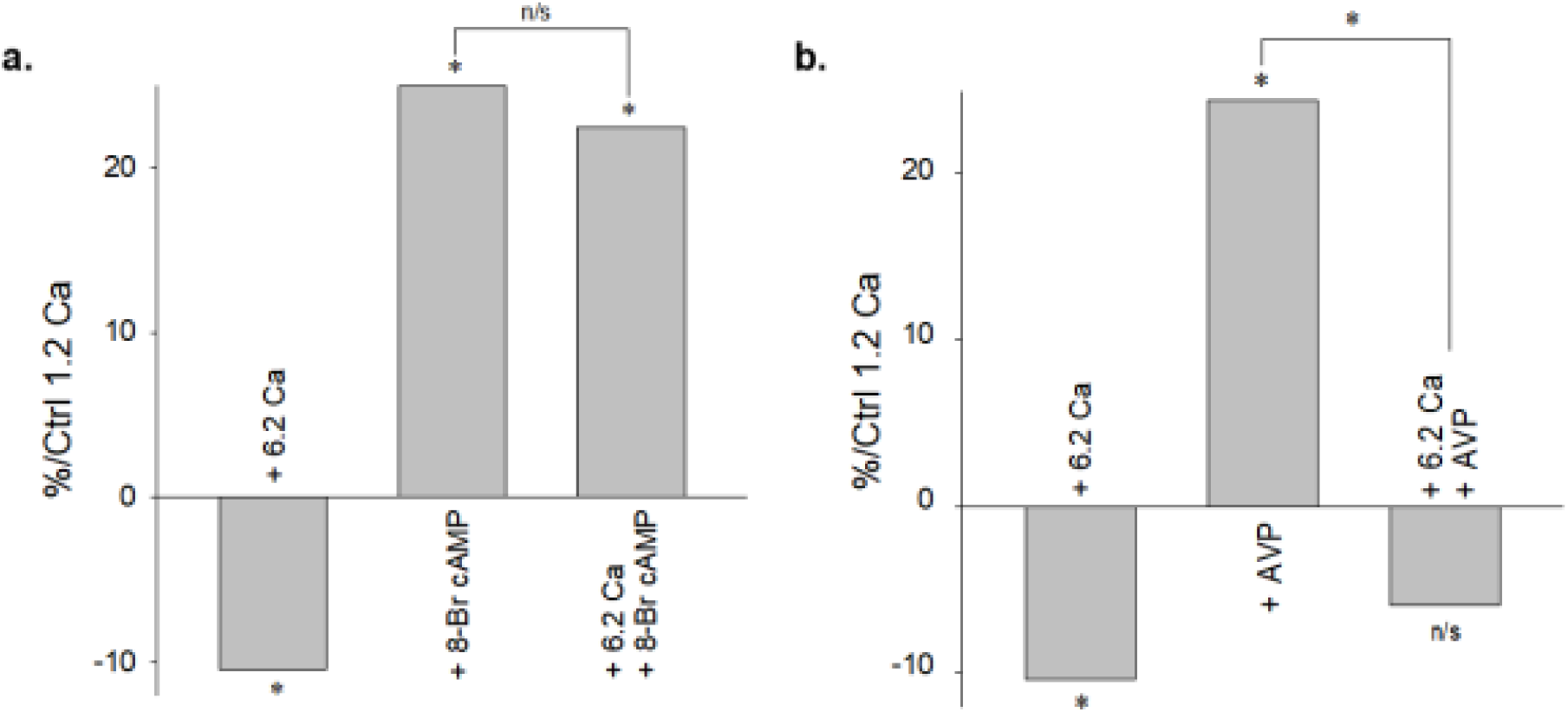
Differences in the length of the primary cilium of LLC-PK1 cells. **a**. Bar graphs represent the percent change either an increase (positive bars) or decrease (negative bars) of the length of the primary cilium of LLC-PK1 cells made relative to normal Ca^2+^ (1.2 mM). Values are shown in the presence either of high Ca^2+^ (+6.2 Ca), 8-Br-cAMP or 6.2 mM Ca^2+^ plus 8-Br-cAMP. **b**. Comparable results obtained in the presence of AVP. Values were made relative to normal Ca^2+^ (1.2 mM). Values are shown in the presence either of high Ca^2+^ (+6.2 Ca), AVP or 6.2 mM Ca^2+^ plus AVP.

### Effect of AVP and high Ca^2+^ on the length of primary cilia in LLC-PK1 cells

To further explore the effect of the cAMP pathway on the length of primary cilia, wild type LLC-PK1 cells were also exposed to AVP (10 µM) under normal external Ca^2+^ conditions (1.2 mM, Fig. 1b). AVP treatment produced a 24.4% increase in primary cilia length (4.38 ± 0.08, n = 314, vs. 5.45 ± 0.09 µm, n = 140, p < 0.0001, Figs. 1b & 2b). A similar experiment conduced in high external Ca^2+^ (6.2 mM), rendered an average ciliary length of 4.12 ± 0.11 µm, n = 120 in the presence of AVP, which only represented a non-significant change in length (5.1%, 0.20 μm length) respect to its control in high Ca^2+^ (3.92 ± 0.11, n = 242, p = 0.2523, Fig. 1a & b). Summarized data shown in Table 2 and in Fig. 2b, indicate that AVP treatment elongated ciliary length, and the effect was much lower in high Ca^2+^ (24.4% vs. 5.1%). This was strictly caused by the shortening effect of the ion. Thus, AVP and external Ca^2+^ effects were independent and additive, such that net effect was a small reduction in length caused by the combined effects of the hormone and high Ca^2+^, which was 5.9% (∼0.26 μm, not-significant, Fig. 1b) shorter as compared to the control condition in normal Ca^2+^ (Table 2).

**Table 2.** Effect of Ca^2+^ and AVP on the length of primary cilia

| | 1.2 Ca<sup>2+</sup>, $\mu\text{m}$ | 6.2 Ca<sup>2+</sup>, $\mu\text{m}$ | $\Delta$, $\mu\text{m}$ | % |
| --- | --- | --- | --- | --- |
| Ctrl, $\mu\text{m}$ | 4.38 ± 0.08<br>n = 314 | 3.92 ± 0.11<br>n = 242 | -0.46<br>(*p* = 0.0006) | -10.5 |
| +AVP, $\mu\text{m}$ | 5.45 ± 0.09<br>n = 140 | 4.12 ± 0.11<br>n = 12 | -1.33<br>(*p* < 0.0001) | -24.4 |
| $\Delta$, $\mu\text{m}$ | 1.07<br>(*p* < 0.0001) | 0.20<br>(*p* = 0.2523) | 0.87 | |
| % | +24.4 | +5.1 |  |  |

### Effect of 8-Br-cAMP on the length of primary cilia from PKD2 gene silenced cells

Polycystin-2, which is functionally present in primary cilia of renal epithelial cells, is regulated by both external Ca^2+^ and the cAMP pathway (Cantero 2013, Cantero 2015, Dai 2017). However, little is known (Perez 2020) as to whether PC2 modifies the length of the primary cilium. Thus, LLC-PK1 cells were treated with PC2-specific (and non-specific) siRNAs to elicit PKD2 gene silencing, as recently reported (Dai 2017, Perez 2020). Two siRNA-treated cell groups were tested, including cells incubated with a non-specific probe (Irss) and cells treated with the probe complementary to PKD2 mRNA (P1ss), which we have recently shown produce an almost complete inhibition of PC2 function in these cells (Dai 2017, Perez 2020). Transfection efficacy was evaluated using a fluorescent silencing probe (data not shown). In the presence of normal Ca^2+^, Irss cells incubated with scrambled probe, were first tested for both 8-Br-cAMP and/or high Ca^2+^ conditions prior to exploring PKD2 silencing conditions. Treatment of Irss cells with 8-Br-cAMP in normal Ca^2+^ produced, in average, a 1.78 μm increase (43%) in ciliary length (5.92 ± 0.15 μm, n = 157, vs. 4.14 ± 0.12 μm, n = 131, p < 0.0001, Fig. 3a). This effect was 60.4% bigger as compared with the effect observed in control cells (1.11 μm, Tables 3 vs. 1).

**Table 3.** Effect of Ca^2+^ and 8-Br-cAMP on the length of primary cilia from Irss-treated cell

| | 1.2 $\text{Ca}^{2+}$, $\mu\text{m}$ | 6.2 $\text{Ca}^{2+}$, $\mu\text{m}$ | $\Delta$, $\mu\text{m}$ | % |
| --- | --- | --- | --- | --- |
| **Ctrl, $\mu\text{m}$** | 4.14 ± 0.12<br>n = 131 | 3.66 ± 0.32<br>n = 45 | **-0.48**<br>($p = 0.0852$) | **-11.6** |
| **+ 8-Br-cAMP, $\mu\text{m}$** | 5.92 ± 0.15<br>n = 157 | 5.43 ± 0.12<br>n = 144 | **-0.49**<br>($p = 0.0122$) | **-8.3** |
| **$\Delta$, $\mu\text{m}$** | **1.78**<br>($p < 0.0001$) | **1.77**<br>($p < 0.0001$) | **0.01** | |
| **%** | **+43.0** | **+48.4** |  |  |

**Fig. 3:**
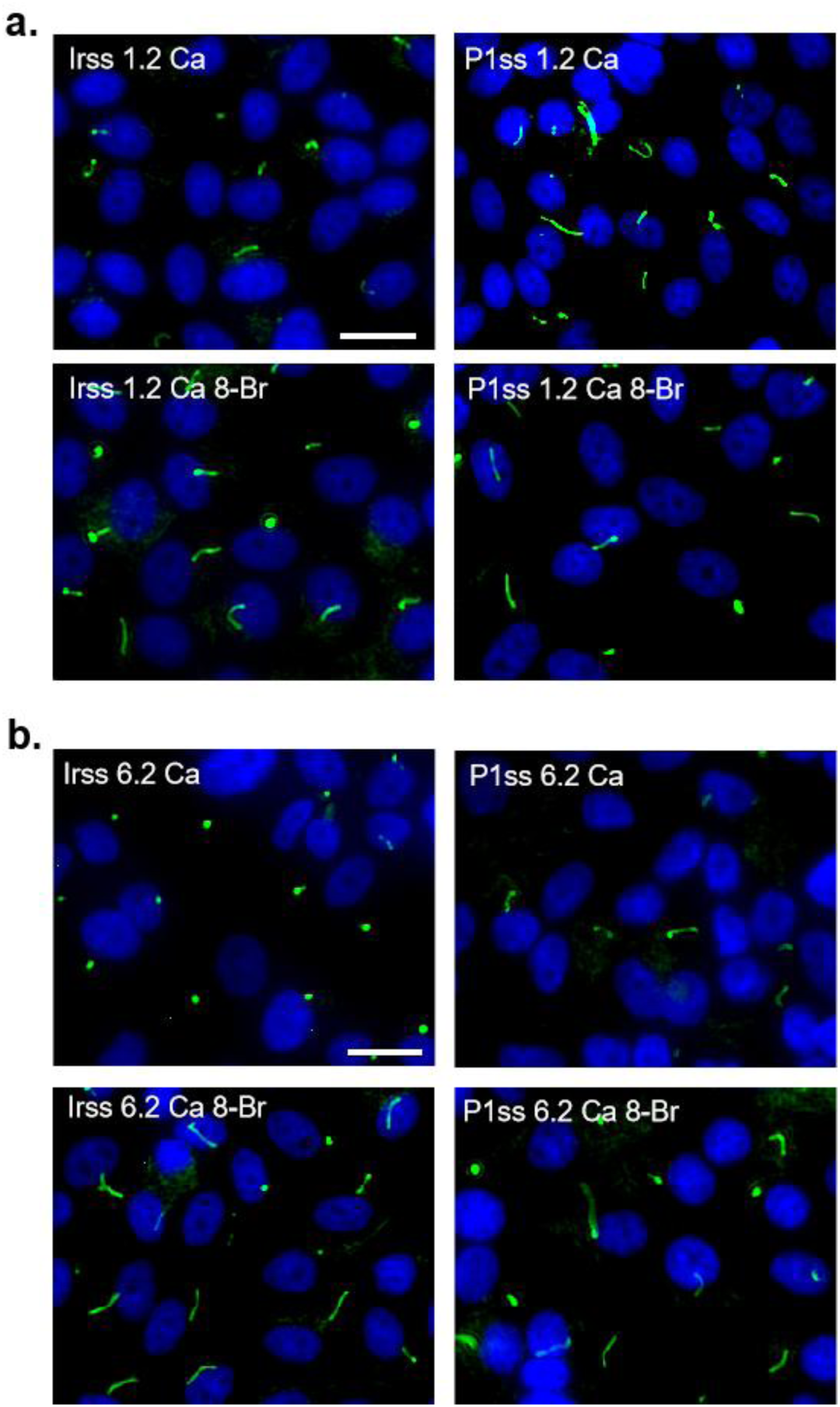
Effect of 8-Br-cAMP on the length of primary cilia from PKD2 gene silenced cells. Confluent monolayers of LLC-PK1 cells were labeled with an anti-acetylated α-tubulin antibody (Green), and DAPI (Blue). The merged images are shown. **a**. In normal Ca^2+^, LLC-PK1 cells were either treated with non-specific (Irss) or PC2-specific (P1ss) siRNAs. 8-Br-cAMP increased the length of primary cilia in Irss-treated cells but not P1ss-treated cells. **b**. In the presence of high Ca^2+^, Irss-treated cells had shorter primary cilia, an effect that disappear after P1ss treatment. 8-Br-cAMP increased primary cilia length after Irss treatment with similar effect on P1ss treated cells.

Irss treatment with lipofectamine to insert the scrambled probe, likely helped delivery of increased 8-Br-cAMP to the cells, thus rendering higher values as compared to control cells (Perez 2020). However, high Ca^2+^ decreased ciliary length by 0.48 μm (11.6%, 4.14 ± 0.12 μm, n = 131, vs. 3.66 ± 0.32, μm, n = 45, p = 0.0852, Fig. 3b & 4b), a finding similar to that observed in control cells (Table 1). Summarized data in Table 3 indicate that in Irss-treated cells 8-Br-cAMP elongated ciliary length to the same extent in both normal and high Ca^2+^ (43.0% vs. 48.4%). The combined 8-Br-cAMP and high Ca^2+^ effects were independent and additive, which rendered a 1.28 μm increase in length (5.43 ± 0.12 μm, n = 144), representing a 31.1% respect to the control value (Fig. 5a, p < 0.0001). With these controls, values were then compared to those of PC2-silenced cells (P1ss). In normal Ca^2+^, P1ss-treated cells had, in average, a statistically significant 1.21 µm increase in length, which represented 29.2% longer primary cilia as compared to Irss-treated cells (4.14 ± 0.12 µm, n = 131 vs. 5.35 ± 0.18 μm, n = 74, p < 0.0001, Figs. 3a & 4a, compare Tables 3 and 4), as recently reported (Perez 2020). P1ss cells were then tested for both 8-Br-cAMP and high Ca^2+^ conditions. In normal Ca^2+^, 8-Br-cAMP treatment of P1ss cells produced a not significant 0.26 μm decrease (4.9%) in ciliary length (5.09 ± 0.10 μm, n = 260, vs. 5.35 ± 0.18 μm, n = 74, p = 0.2179). However, high Ca^2+^ alone induced a statistically significant 0.94 μm decrease (17.6%) in ciliary length (4.41 ± 0.34 μm, n = 37, vs. 5.35 ± 0.18 μm, n = 74, p = 0.0083). The combined effect of 8-Br-cAMP and high Ca^2+^, rendered a 0.92 μm increase (20%) in length (5.33 ± 0.12 μm, n = 129, p = 0.0017, Fig. 3b & 4b). Paradoxically, the combined maneuvers returned the values to control values (5.33 ± 0.12 µm, n = 129 vs. 5.35 ± 0.18 µm, n = 74, Fig. 5b, Tables 4, A1 & A2).

**Table 4.** Effect of Ca^2+^ and 8-Br-cAMP on the length of primary cilia from P1ss-treated cells

| | 1.2 $\text{Ca}^{2+}$, $\mu\text{m}$ | 6.2 $\text{Ca}^{2+}$, $\mu\text{m}$ | $\Delta$, $\mu\text{m}$ | % |
| --- | --- | --- | --- | --- |
| Ctrl, $\mu\text{m}$ | 5.35 ± 0.18<br>n = 74 | 4.41 ± 0.34<br>n = 37 | -0.94<br>(p = 0.0083) | -17.6 |
| + 8-Br-cAMP, $\mu\text{m}$ | 5.09 ± 0.10<br>n = 260 | 5.33 ± 0.12<br>n = 129 | 0.24<br>(p = 0.1472) | +0.5 |
| $\Delta$, $\mu\text{m}$ | -0.26<br>(p = 0.2179) | 0.92<br>(p = 0.0017) | 0.7 | |
| % | +4.9 | +20 |  |  |

**Fig. 4:**
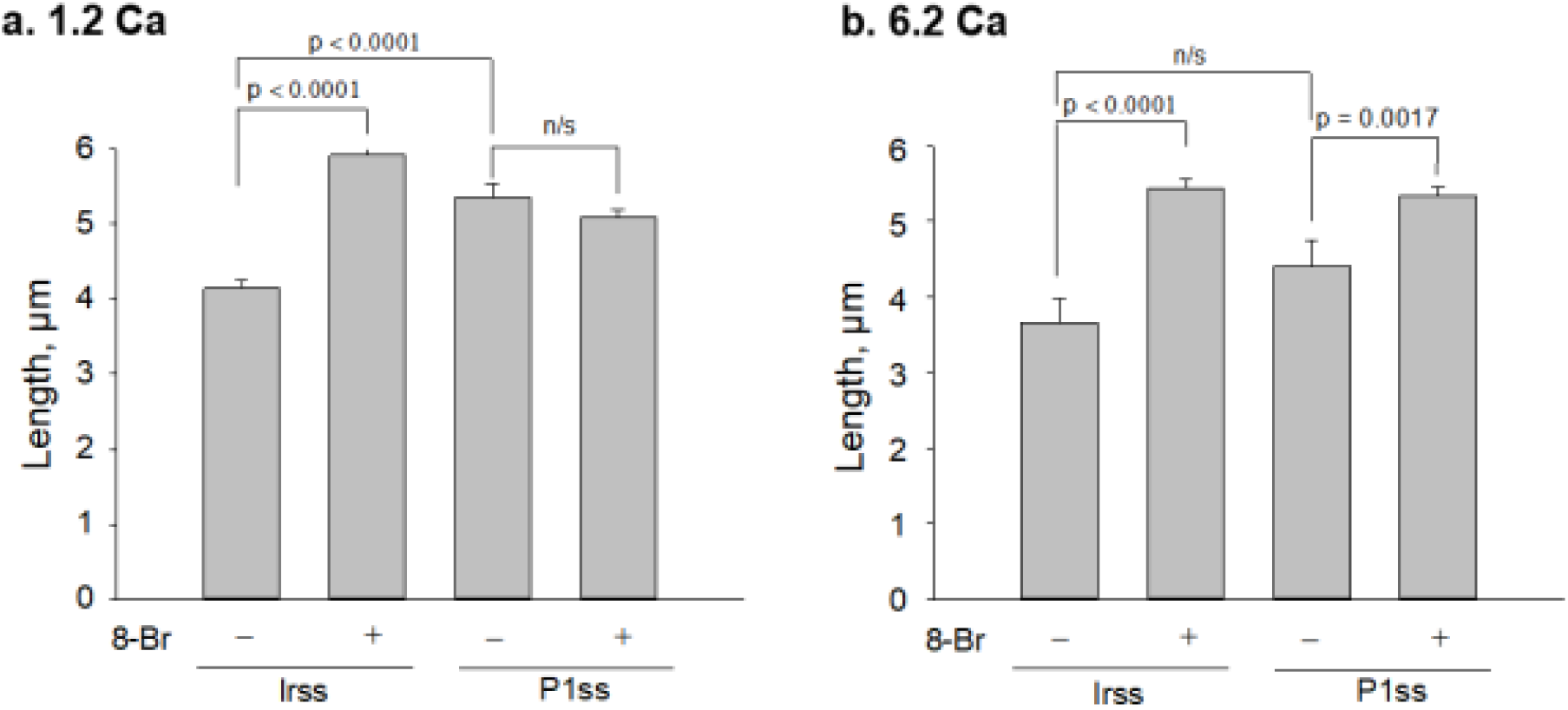
Cilia length comparisons of Irss- and P1ss-treated LLCPK1 cells. Bar graphs represent Mean ± SEM of primary cilium length either in the presence (+) or absence (-) of 8-Br-cAMP for Irss and P1ss treated cells, in the presence of either 1.2 mM Ca^2+^ (**a**.) or 6.2 mM Ca^2+^ (**b**.). In normal Ca^2+^, 8-Br-cAMP failed to modify ciliary length in P1ss-treated cells. However, this effect was partially reversed in high Ca^2+^ (b). Statistical differences are shown as indicated.

**Fig. 5:**
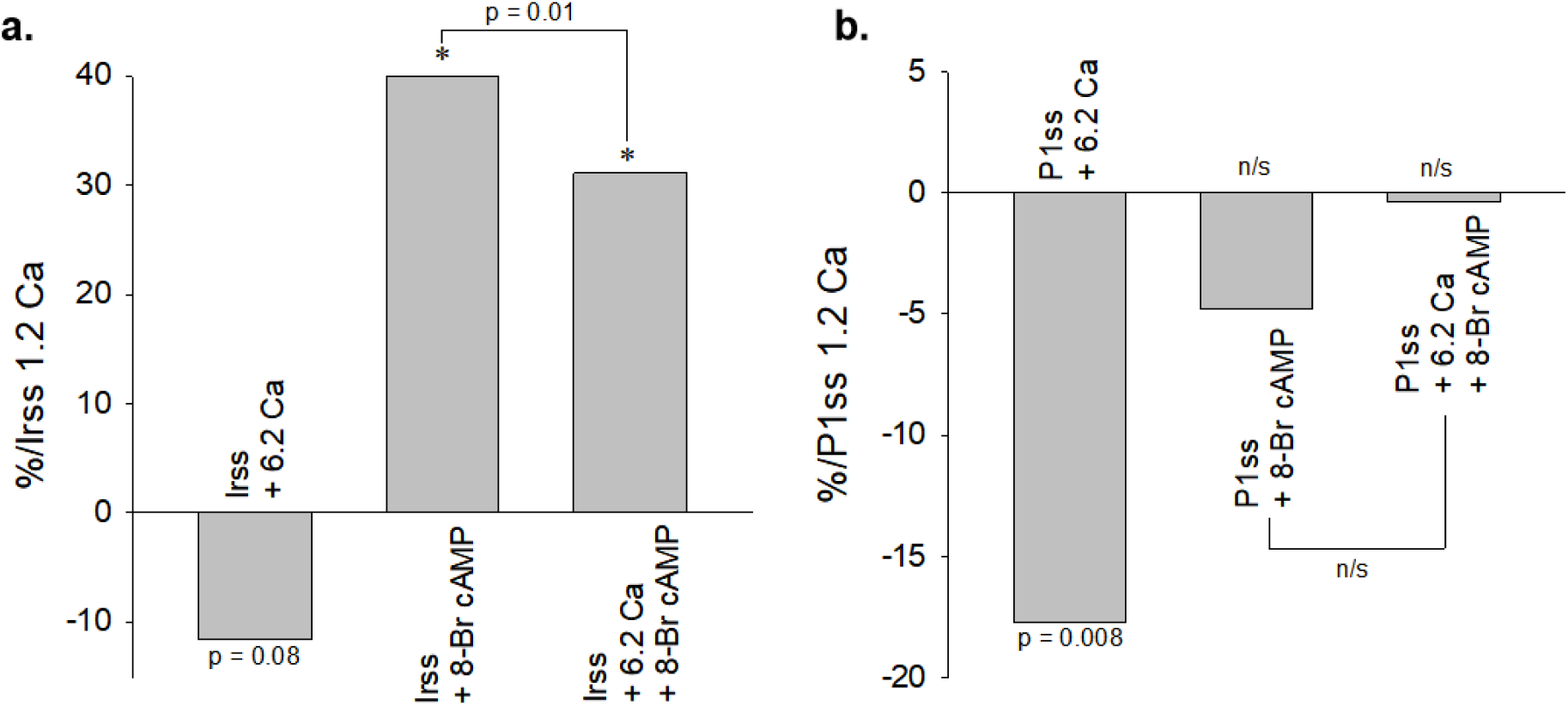
Changes in ciliary length by Ca^2+^ and 8-Br-cAMP in PC2 silenced cells. Bar graphs represent percent changes under different conditions respect to either Irss (**a**.) or P1ss (**b**.) cells in normal Ca^2+^, respectively. **a**. 8-Br-cAMP induced an increase in ciliary length in both normal and high Ca^2+^. **b**. High Ca^2+^ induced a severe shortening of primary cilium, which was entirely reversed by 8-Br-cAMP.

## Discussion

The primary cilia of renal epithelial cells are thought to be mechanosensors implicated in the transduction of environmental signals, which elicit Ca^2+^ entry steps, and cell activation (Praetorius 2001, Nauli 2003, Praetorius 2003, Bergmann 2008, Streets 2013). The structure, function and stability of primary cilia are all essential in normal kidney development. Conversely, cilia dysfunction, including abnormalities in cilia structure, composition, and signaling (Wheatley 1995, Hildebrandt 2007, Watnick 2003) have been linked to various genetic disorders. A number of mutated proteins associated with renal cystic diseases localize to either the primary cilium or its associated structures, including the basal body, centrosomes or ciliary transition zone (Hildebrandt 2005). There are more than a hundred known ciliopathies, including syndromes such as Bardet-Biedl (BBS), Joubert (JBTS), Meckel-Gruber (MKS), Alström, Senior-Löken, and Oro-facial-digital type 1 (OFD1), and diseases such as autosomal dominant polycystic kidney disease (ADPKD) and nephronophthisis (NPHP) (Wheatley 1995, Hildebrandt 2007). However, comparatively little is known as to how primary cilium dysfunction may explain the wide variety of renal phenotypes seen in the different diseases (Loftus 2013). A relevant link between primary cilia and renal cyst formation in ADPKD has been the early finding that PC1 and PC2 colocalize to the primary cilium (Yoder 2002, Ong 2013). However, ciliary length seems to be normal in cells and organisms carrying *PKD1* and *PKD2* mutations, thus implying functional defects rather than structural ones (Ong 2013).

It is currently accepted that in primary cilia of renal epithelia, PC1 and PC2 form a complex that initiates mechanosensory Ca^2+^ signals that control epithelial cell behavior (Nauli 2003). Mechanistically, this is associated with the properties of PC2, itself a Ca^2+^-permeable cation channel implicated in Ca^2+^ signals (González-Perrett 2001). However, how this channel activity may be linked to functional/structural correlated of ciliary parameters still is a matter of study. One relevant structural parameter of primary cilia is its length, which is modified under various physiological and pathological conditions. Murine-injured kidneys (Mahuzier 2012, Simons 2005, Pugacheva 2007) and human renal transplants suffering from acute tubular necrosis (Pugacheva 2007) show longer primary cilia than normal. Toxic Cobalt chloride-treated renal epithelial cells also show much longer cilia than control cells. Thus, it is speculated that primary cilia elongation following renal injury may actually increase their sensory capacity, which is crucial to epithelial differentiation during renal repair. Lithium, which is a potent pharmacological agent (Machado-Vieira 2009), lengthens primary cilia in the mouse brain and cultured cells, including neurons and fibroblasts (Miyoshi 2009). Taken together these findings with the fact that PC2 localizes to renal primary cilia (Nauli 2003, Raychowdhury 2005), and that Li^+^ inhibits its function in vitro (Cantero 2011), it is tempting to link ciliary channel dysfunction to changes in primary cilia parameters. A recent study from our laboratory explored the effect(s) of PC2 channel blockers and *PKD2* gene silencing on the length of primary cilia in wild type LLC-PK1 renal epithelial cells (Perez 2020). Both amiloride and Li^+^, which inhibit PC2 channel function, significantly increased primary ciliary length. This phenomenon was mimicked by specific siRNA *PKD2* gene silencing. In contrast, it was observed that, cells exposed to a high external Ca^2+^ concentration had shorter primary cilia as compared to their controls in normal Ca^2+^. Thus, maneuvers that prevent PC2 function lengthen primary cilia while increased Ca^2+^ influx to the organelle likely collapses axoneme growth, rendering shorter cilia.

The present study was conducted to assess potential regulatory connections between Ca^2+^ and cAMP. These two universal messengers crosstalk, such that a rise in one sets feedback mechanisms to lower the other one and vice versa (Bell 1985, Bugrim 1999, Berridge 2000, Borodinsky 2006, Bruce 2003, Siso-Nadal 2009). Herein we explored whether cAMP-associated signals contributed to the regulation of the length of the primary cilium in LLC-PK1 renal epithelial cells and whether PC2 could be implicated in this regulation. We observed that exposure to 8-Br-cAMP in normal Ca^2+^ lengthened the primary cilium, a phenomenon that was mimicked by treatment with AVP, which increases intracellular cAMP in both renal epithelia and LLC-PK1 cells (Brown 2000, Bouley 2005, Raychowdhury 2009). Exposure of control cells to high Ca^2+^, in contrast, shortened primary cilia length, as recently reported (Perez 2020). The combined effect of Ca^2+^ and 8-Br-cAMP, lengthened the primary cilium, a phenomenon not shared by the combined effects of high Ca^2+^ and AVP, which instead rendered a smaller, but significant shortening in the length of the primary cilium. Thus, the 8-Br-cAMP and AVP maneuvers had opposite effects in high Ca^2+^.

The anti-diuretic regulatory influence of AVP on transepithelial transport is mediated by V2R-cAMP responses triggered from the basolateral side of target epithelia (Brown 2000). However, AVP also has luminal effects associated with both, V2R (Fenton 2007) and Type 1, V1R receptors, which are associated with a rise in Ca^2+^ (Wong 1988, Weinberg 1989, Burgess 1994). Thus, the differences between AVP and 8-Br-cAMP on primary cilia exposed to high Ca^2+^ could be linked to the differential activation of vasopressin receptors which, in turn, may be linked to opposite downstream Ca^2+^ and cAMP signals. Conversely, competition between different receptors such as V2R and CaSR (Dai 2017) may elicit a compensatory response. Ca^2+^/cAMP signaling implicate different targets including specific adenylyl cyclases (ACs) and phosphodiesterases that either increase or blunt cAMP production, respectively. We previously showed a functional coupling between V2R and type V/VI adenylyl cyclase. The entire cAMP pathway induced the production of cAMP in primary cilia upon AVP treatment (Raychowdhury 2009). The regulation by cAMP of cilia structure may be of particular interest because evidence has accumulated for cAMP to play an important role in the cystogenesis of ADPKD (Grantham 2003). Endogenous AC agonists have a stimulatory effect on both cell proliferation and fluid transport in various forms of cystic renal epithelia (Belibi 2004), and ADPKD occurs with high levels of cAMP and low levels of Ca^2+^ (Yamaguchi 2000, Starremans 2008). Recent evidence indicated that AVP antagonists are effective in ameliorating cystic disease (Torres 2004, Wang 2005, Torres 2014).

To assess whether PC2 was implicated in the role the cAMP pathway had on primary cilia, the PKD2 gene product was silenced, as recently reported (Dai 2017, Perez 2020). In normal Ca^2+^, PC2-silenced cells had statistically longer primary cilia by as much as 1.22 µm (22.61%) compared to their own controls (Irss-treated cells). However, 8-Br-cAMP treatment of the PC2-silenced cells showed a very small, 0.26 µm increase (4.86%) in ciliary length, as compared to the dramatic 43% increase in ciliary length (1.78 µm) in their respective controls (Irss-treated cells). The combined effect of both second messengers rendered a 1.46 μm (45.6%) lengthening as compared to the Irss-treated cells, 0.95 μm (22.9%). Thus, 8-Br-cAMP plus high Ca^2+^ elicited the largest lengthening of primary cilia in the absence of PC2, suggesting regulatory mechanisms of ciliary structure other than those associated with the channel in the control of this parameter. The presence of V2R and a functional cAMP pathway in primary cilia (Raychowdhury 2009) would contribute to cAMP-mediated signals and the balance of Ca^2+^ homeostasis (Bell 1985, Gao 1999, Merritt 1992), which in turn would help collapse axoneme microtubules (Karr 1980, O’Brien 1997), shortening primary cilia. PC2 function itself in primary cilia is regulated by axoneme structures in renal epithelial cells (Li 2006).

In summary, we provide evidence for a regulatory role of the cAMP pathway in the control of primary cilia length in renal epithelial cells. Both, the presence of V2R and a functional cAMP pathway contributed to this phenomenon, albeit by different signaling mechanisms. This, in turn suggests that in combination with, or in addition to, PC2 channel function, receptor-mediated second messenger pathways, may be key signaling components in the sensory properties of primary cilia. Both Ca^2+^ and cAMP remain two different branches of a combined signaling pathway that may regulate similar target effector systems. Our results further suggest that PC2 plays a role in this regulation. Maneuvers leading to its inhibition would render ciliary elongation as a consequence of a decrease in ciliary Ca^2+^ entry. This mechanism is under the control of cAMP signals. Our results further suggest that both second messengers may act in more complicated ways than previously expected, such that the relationship between them may lead to changes in ciliary length and the formation of renal cysts in ADPKD.

## Notes

### Competing Interest Statement

The authors have declared no competing interest.

