## Appendix for "Ca^2+^-Dependent Regulation by the Cyclic AMP Pathway of Primary Cilium Length in LLC-PK1 Renal Epithelial Cells"

**Table 1A.** Effect of PC2 silencing and 8-Br-cAMP on the length of primary cilium in the presence of 1.2 mM $Ca^{2+}$

| | Irss cells, µm | P1ss cells, µm | Δ, µm | % |
|---|---|---|---|---|
| **Ctrl, µm** | 4.14 ± 0.12<br>n = 131 | 5.35 ± 0.18<br>n = 74 | **1.21**<br>**(*p* < 0.0001)** | **+29.2** |
| **+ 8-Br-cAMP, µm** | 5.92 ± 0.15<br>n = 157 | 5.09 ± 0.10<br>n = 260 | **-0.83**<br>**(*p* < 0.0001)** | **+14.0** |
| **Δ, µm** | **1.78**<br>**(*p* < 0.0001)** | **-0.26**<br>**(*p* = 0.2179)** | **0.38**<br>**1.52** | |
| **%** | **+42.0** | **-4.9** | | |

**Table 2A.** Effect of silencing and 8-Br-cAMP on the length of primary cilium in the presence of 6.2 mM $Ca^{2+}$

| | Irss cells, µm | P1ss cells, µm | Δ, µm | % |
|---|---|---|---|---|
| **Ctrl, µm** | 3.66 ± 0.32<br>n = 45 | 4.41 ± 0.34<br>n = 37 | **0.75**<br>**(*p* = 0.1134)** | **+20.5** |
| **+ 8-Br-cAMP, µm** | 5.43 ± 0.12<br>n = 144 | 5.33 ± 0.12<br>n = 129 | **-0.1**<br>**(*p* = 0.5574)** | **-1.8** |
| **Δ, µm** | **1.77**<br>**(*p* <0.0001)** | **0.92**<br>**(*p* = 0.0017)** | **0.65**<br>**1.68** | |
| **%** | **+48.4** | **+20.9** | | |

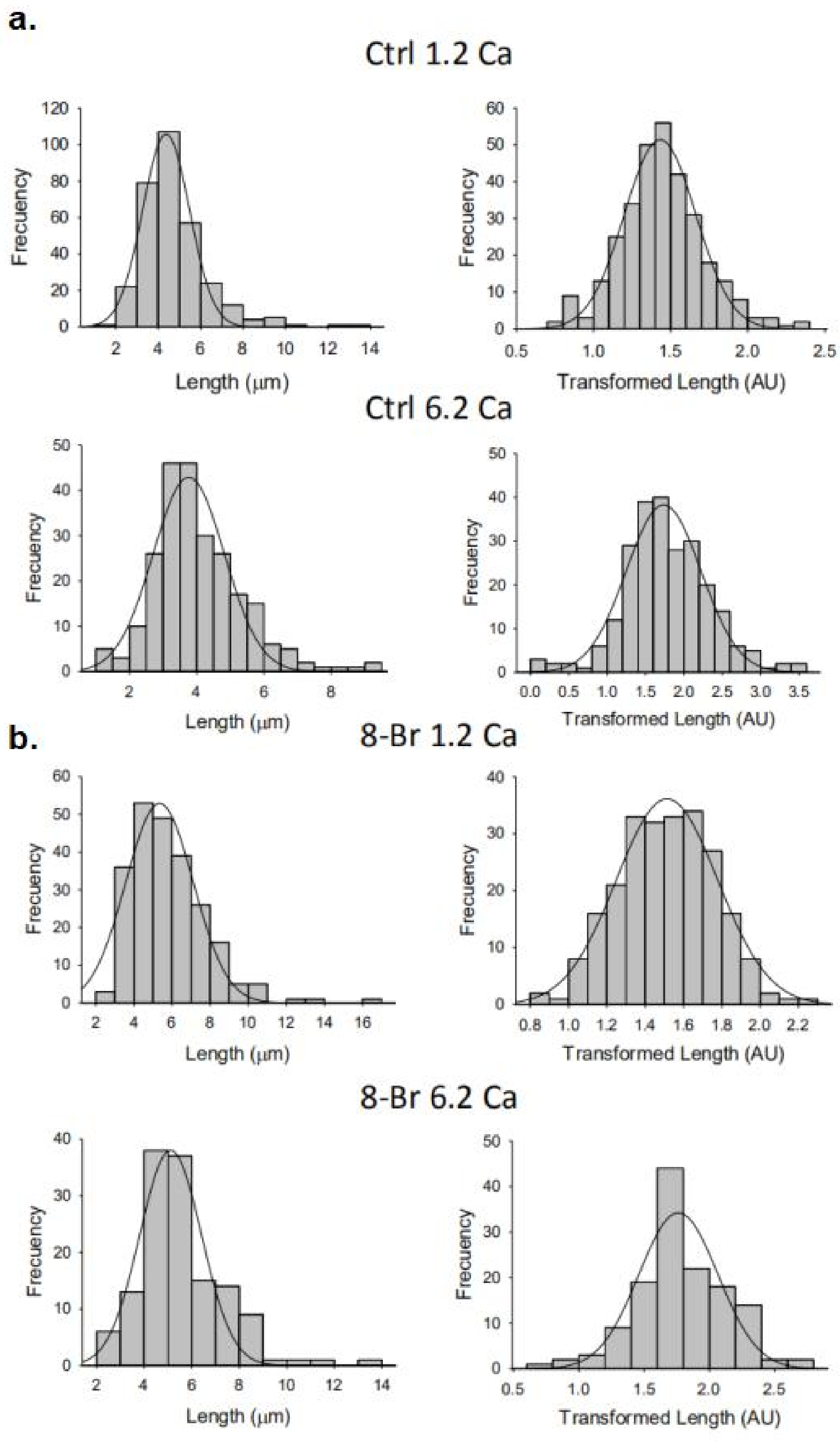


***Fig. 1A.*** *Ciliary length distribution in the presence of $Ca^{2+}$ and 8-Br-cAMP.* **a.** Histograms of ciliary length measurements in normal (1.2 mM) and high (6.2 mM) $Ca^{2+}$ are shown before (Left), and after (Right), Box-Cox transformation. A not Normal left-skewed distribution of data is observed before transformation. **b.** Histograms of ciliary length measurements in the presence of 8-Br-cAMP and either normal (8-Br 1.2 mM) or high (8-Br 6.2 mM) $Ca^{2+}$ are shown before (Left), and after (Right), Box-Cox transformation. A not Normal left-skewed distribution of data is observed before transformation.

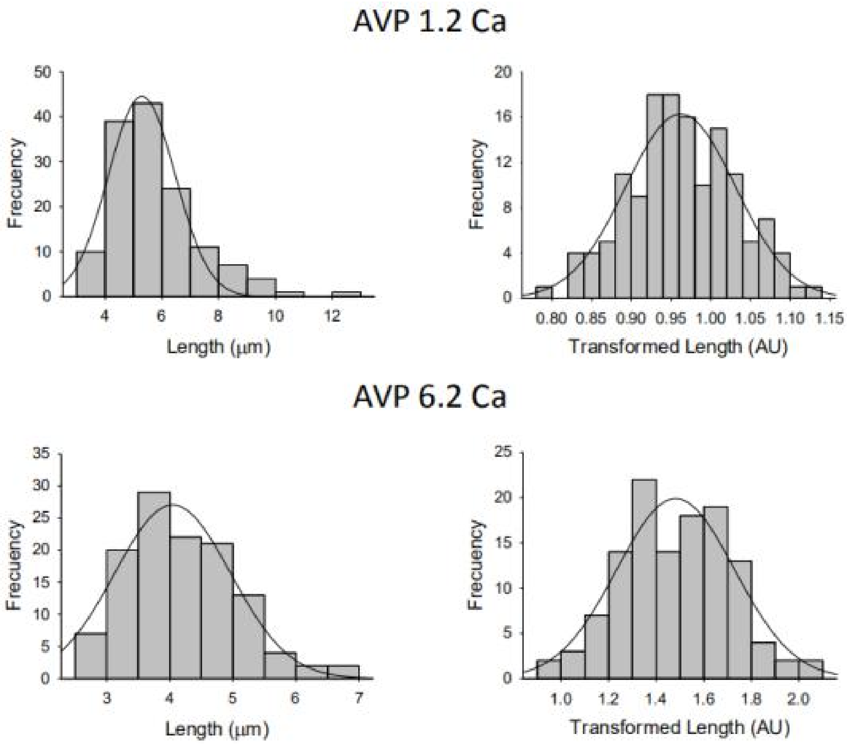


***Fig. 2A.*** *Ciliary length distribution in the presence of $Ca^{2+}$ and AVP.* **a.** Histograms of ciliary length measurements in the presecne of AVP and either normal (AVP 1.2 mM) or high (AVP 6.2 mM) $Ca^{2+}$ are shown before (Left), and after (Right), Box-Cox transformation. A not Normal left-skewed distribution of data is observed before transformation.

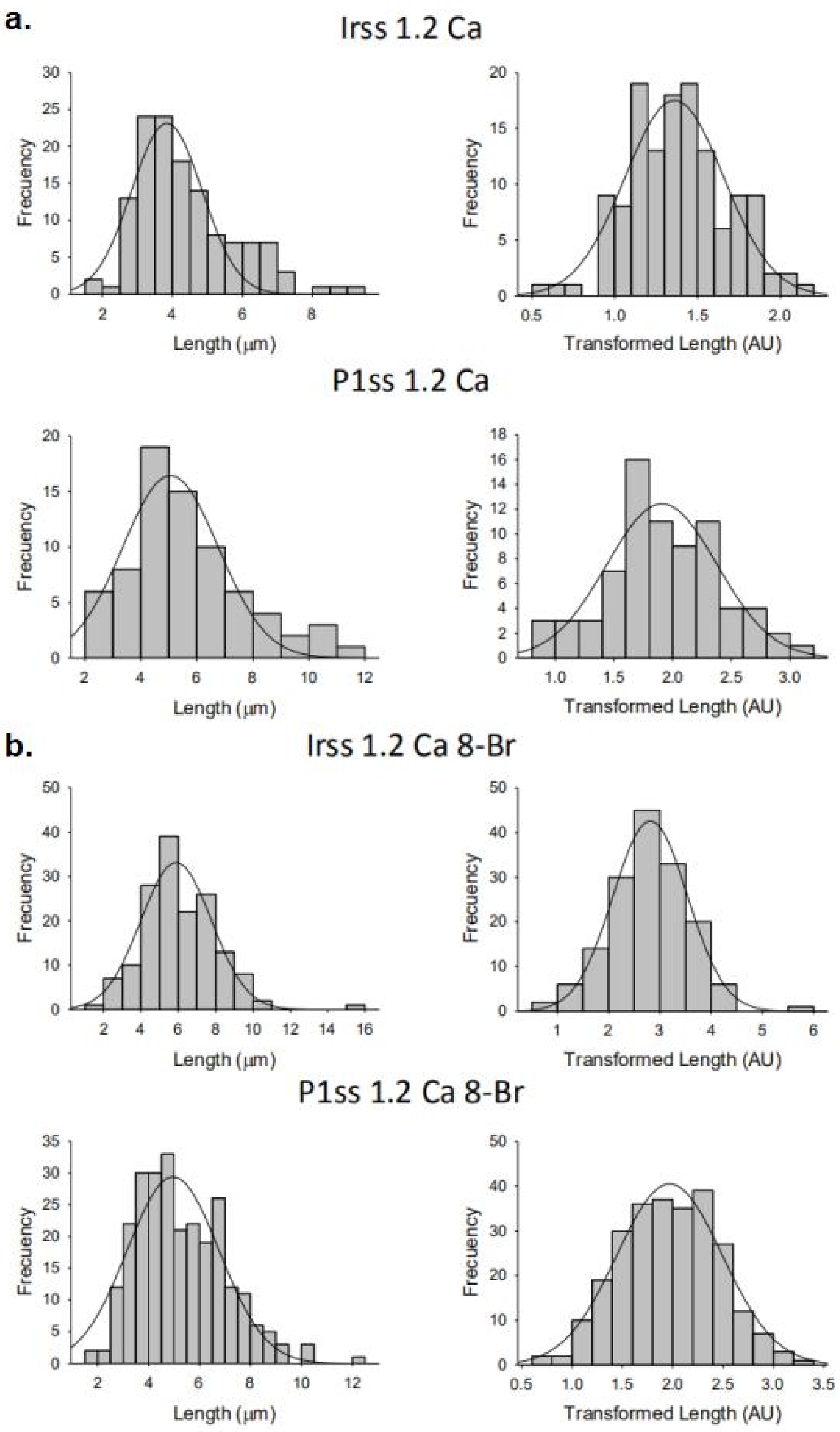


***Fig. 3A.*** *Ciliary length distribution in the presence of 1.2 mM $Ca^{2+}$ and 8-Br-cAMP of either Irss or P1ss-teated cells.* **a.** Histograms of ciliary length measurements in normal $Ca^{2+}$ of either Irss-treated (Irss 1.2 Ca) or P1ss-treated (P1ss 1.2 Ca) cells are shown before (Left), and after (Right), Box-Cox transformation. A not Normal left-skewed distribution of data is observed. b**.** Histograms of ciliary length measurements in normal $Ca^{2+}$ and 8-Br-cAMP of either Irss-treated (Irss 1.2 Ca 8-Br) or P1ss-treated (P1ss 1.2 Ca 8-Br) cells are shown before (Left), and after (Right), Box-Cox transformation. A not Normal left-skewed distribution of data is observed before transformation.

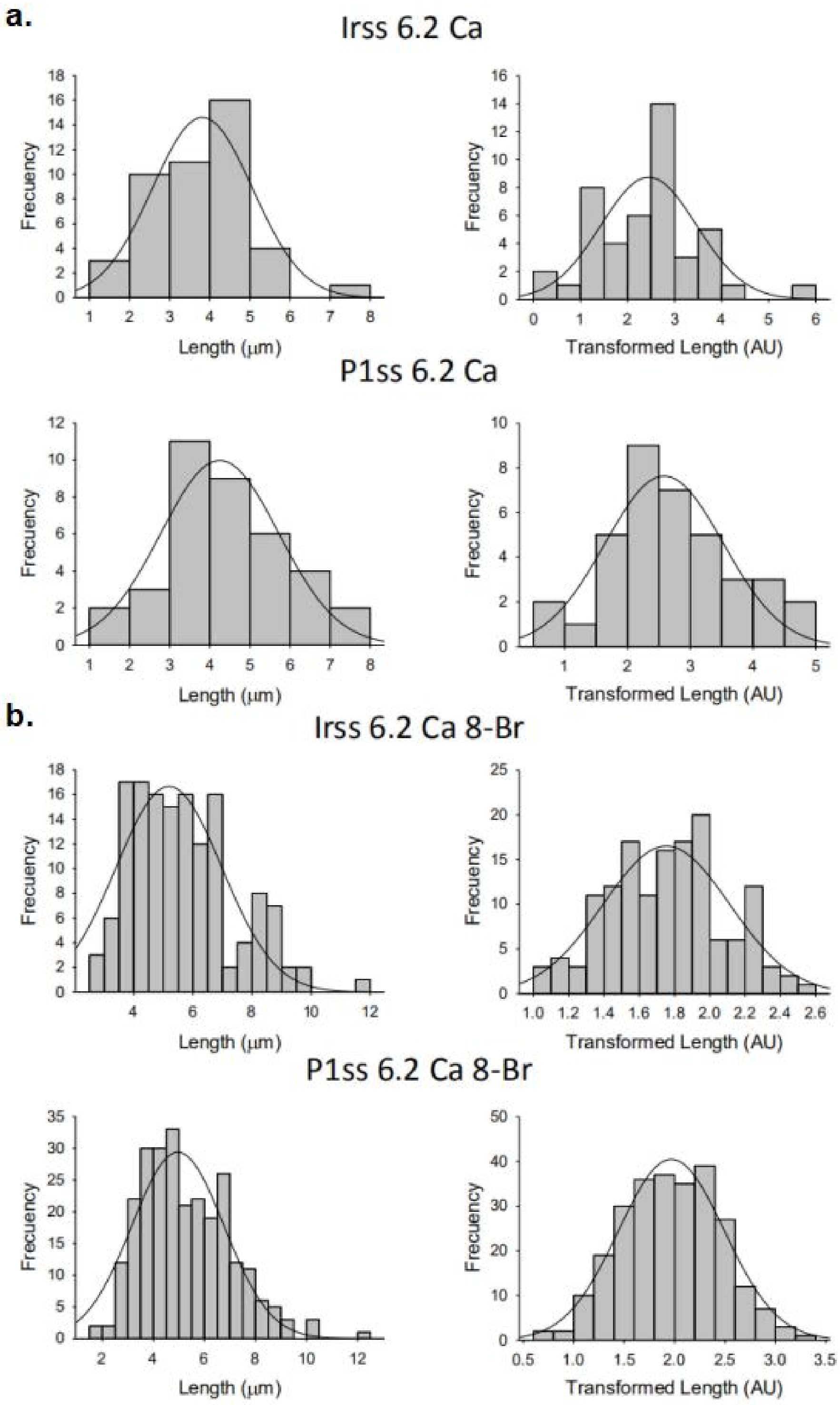


***Fig. 4A.*** *Ciliary length distribution in the presence of 6.2 mM Ca$^{2+}$ and 8-Br-cAMP of either Irss or P1ss-teated cells.* **a.** Histograms of ciliary length measurements in high Ca$^{2+}$ of either Irss-treated (Irss 6.2 Ca) or P1ss-treated (P1ss 6.2 Ca) cells are shown before (Left), and after (Right), Box-Cox transformation. A not Normal left-skewed distribution of data is observed. b**.** Histograms of ciliary length measurements in normal Ca$^{2+}$ and 8-Br-cAMP of either Irss-treated (Irss 6.2 Ca 8-Br) or P1ss-treated (P1ss 6.2 Ca 8-Br) cells are shown before (Left), and after (Right), Box-Cox transformation. A not Normal left-skewed distribution of data is observed before transformation.
